# AtHDA15 attenuates COP1 *via* transcriptional quiescence, direct binding, and sub-compartmentalization during photomorphogenesis

**DOI:** 10.1101/2020.08.07.241737

**Authors:** Malona V. Alinsug, Custer C. Deocaris

## Abstract

Light is an essential environmental cue that determines the overall growth and development of plants. However, the molecular mechanisms underpinning the light signaling network are obscured by the epigenetic machinery where reversible acetylation and deacetylation play crucial roles in modulating light-regulated gene expression. In this paper, we demonstrate that HDA15 represses COP1, the master switch in the light signaling network, by deacetylation, protein interaction, and sub-compartmentalization. *hda15* T-DNA mutant lines exhibited light hyposensitivity with significantly reduced HY5 and PIF3 transcript levels leading to long-hypocotyl phenotypes in the dark while its overexpression exhibited elevated HY5 transcripts and short hypocotyl phenotypes. *In vivo* and *in vitro* binding assays further show that HDA15 directly interacts with COP1 inside the nucleus modulating COP1’s repressive activities. Crossing *hda15-t*27 with *cop1-4* mutants resulted in short-hypocotyl and dwarfed phenotypes, reminiscent of *cop1-4* mutants suggesting COP1 is epistatic to HDA15. Although light signals the nucleocytoplasmic shuttling of HDA15, the presence of COP1 triggers its nuclear localization. A working model is presented elucidating the concerted interplay between HDA15 and COP1 under light and dark conditions.

## INTRODUCTION

Plants adapt extensively to the dynamic changes in their light environment as they are critically dependent on light for photosynthesis, growth, and development. In general, light induces massive changes in a plant’s transcriptional activity resulting in distinct expression profiles during seedling development (Jiao et al., 2007; Ma et al., 2013). Approximately 15% of these transcriptional changes have been attributed to histone modifications, specifically the acetylation in H3K9K27, which coincides with the upregulation of key photomorphogenesis gene expression, such as HY5 and its homolog, HYH (Charron et al., 2009). H3K9ac is one of the most characterized epigenetic marks invariably correlated with transcriptional activation in all the species studied (Ausin et al., 2004; Kurdistani et al., 2004; Schübeler et al., 2004; Benhamed et al., 2008; Guo et al., 2008; Charron et al., 2009; Zhou et al., 2009; Schenke et al., 2014; Liu et al., 2017). However, most of these light-induced pathways remain obscure, while little is understood about histone deacetylases' involvement in regulating these genes.

Studies on histone active marks such as H3K9K27ac have been limited to positive photomorphogenic regulators such as CABs, LHBIB1, LHCA1, HYH and HY5 and its downstream targets CHS, IAA3, RBCS-1a (Benhamed et al., 2006; Guo et al., 2008; Charron et al., 2009). Among the 12 members of the RPD3/HDA1 superfamily in *Arabidopsis*, HDA19, HDA6, and HDA15 have been ascribed active roles in light-regulated gene expression. Based on studies, HDA19 and HDA6 play a negative regulatory role by repressing some of these light-regulated genes (Benhamed et al., 2006; Tessadori et al., 2009). On the other hand, HDA15 has recently been shown to have a regulatory role in photomorphogenesis. As human Class II HDACs are known to undergo nucleocytoplasmic shuttling, we have demonstrated that HDA15 shuttles to the nucleus under light (Alinsug et al., 2012). Moreover, phytochrome interacting factor 3 (PIF3) associates with HDA15 repressing chlorophyll biosynthesis and photosynthesis in etiolated seedlings (Liu et al., 2013). Similarly, PIF1 recruits HDA15 at the promoter sites repressing the transcription of light-responsive genes involved in multiple hormonal signaling pathways and cellular processes in germinating seeds in the dark (Gu et al., 2017). Furthermore, HDA15 forms a complex with nuclear factor homologs (NF-YC) functioning as a transcriptional co-repressor via deacetylation inhibiting hypocotyl elongation in photomorphogenesis (Tang et al., 2017). A recent study describes that long hypocotyl 5 (HY5) protein interacts with HDA15 repressing hypocotyl cell elongation during photomorphogenesis (Zhao et al., 2019).

HY5 and PIFs are key transcription factors playing crucial regulatory roles in photomorphogenesis and skotomorphogenesis, respectively. Along with the light signaling network, both HY5 and PIFs are repressed by COP1, an E3 ubiquitin ligase that targets both photoreceptors HY5 and HYH for ubiquitination and proteasome degradation in the dark (Osterlund et al., 2000). However, the molecular interplay between COP1, HY5, PIFs, and HDA15 remains unclear. It remains obscure as to how or when HDA15 precisely interjects within the light response molecular network. As epigenetic regulation further adds complexity to the light signaling pathway, we attempt to explore the multiple roles HDA15 plays during photomorphogenesis. In this paper, we demonstrate that HDA15 represses COP1, the master switch in the light signaling network, by transcriptional silencing via deacetylation, protein-protein association, and sub-compartmentalization. A working model is presented elucidating the interplay of the multiple actions exerted by HDA15 on COP1 under light and dark conditions.

## Materials & Methods

### Plant Material

Seeds of T-DNA knockout lines *hda5-t24* (SALK_030624)*, hda15-t27* (SALK_004027C), and *hda18-t* (SALK_006938C) in Col-0 background were purchased from ABRC and screened via PCR using specific primers. Homozygous mutant plants with positive T-DNA inserts were grown under long day and assessed for the expression of the corresponding gene compared to wild type via RT-PCR using specifically designed primers. Seeds of homozygous T-DNA knockout lines were then collected and further used for this study. To establish the double mutant line, we crossed the homozygous *hda15-t27* mutant with the *cop1-4* mutant. The resultant F2 lines were then finally screened by PCR.

### Protoplast PEG Transfection

Leaves of 3-week old Col-0 wild type, T2 transgenic lines, and mutant lines *phyA, phyB, cry1/cry2,* and *cop1-4* were used for protoplast transient expression assay. Isolation of Arabidopsis mesophyll protoplasts and PEG transfection were done as described previously (Yoo et al., 2007) with some modifications. Mesophyll protoplasts were isolated from 3-week old Col-0 plants and Arabidopsis PSB-D cell lines. Twenty μg of HDA15-YFP/GFP fusion plasmid and VirD2-NLS, a nuclear marker, were co-transfected into protoplasts using polyethylene glycol (PEG) solution (0.4 g/ml PEG 4000, 0.8 M mannitol, 125 mM CaCl2). After incubation for 5-15 min at room temperature, the protoplasts were washed and resuspended in W5 solution (154 mM NaCl, 125 mM CaCl2, 5 mM KCl, 2 mM MES at pH5.7), and further incubated under white light for 16–24 h before imaging using Leica SP5 confocal microscope. For the localization of HDA15 in different light treatments, transfected protoplasts were incubated under white light for 18 h then transferred to E30LEDL3 growth chambers (Percival Scientific) with far-red, red, and blue light-emitting diode sources for 3 h. Low light intensities used as treatment were measured at 2.77 μmol m^−2^ s^−2^ (FR), 1.77 μmol m^−2^ s^−2^ (R), and 3.84 μmol m^−2^ s^−2^ (B).

### Light treatments and hypocotyl measurements

Arabidopsis seeds were surface sterilized, cold treated for 3-4 days and sown on 1/2 MS plates with 0.1% sucrose then subjected to different light treatments at 21°C for 5-7 consecutive days. White light intensity was approximately 130 μmol m^−2^s^−1^. Plated seeds were incubated in E30LEDL3 colored light chambers (Percival Scientific) with the corresponding low (L) and high (H) wavelength measurements as follows: far-red at 18.065 (H) and 2.77 (L) μmol m^−2^ s^−1^, red light at 22.938 (H) and 1.77 (L) μmol m^−2^ s^−1^, and blue light at 11.095 (H) and 3.84 (L) μmol m^−2^ s^−1^, respectively. On the other hand, dark treated seedlings were wrapped in foil and placed inside a dark growth chamber. Hypocotyl length was measured using UTHSCA Image Tool using 30-50 seedlings for each line and further analyzed for statistical tests.

### Total RNA extraction & qRT-PCR

Gene expression was initially assessed by semi-quantitative RT-PCR. Total RNA was extracted from plant samples weighing 0.25 to 0.3 g using TRIZOL reagent (Invitrogen). Oligo(dT) primed reverse transcription of first-strand cDNA synthesis was carried out with 7 mg total RNA using Super ScriptTM III (Invitrogen). Equal volumes of each first-strand reaction were amplified with gene-specific primer pairs. Thermocycling conditions were 95°C for 4 mins followed by 30 cycles of 95°C for 30 s, 55–60°C for 30 s, and 74°C for 1–2 min. cDNAs obtained from RT were then used as templates to run real-time PCR. The following components were added to a reaction tube: 9 μL of iQ SYBR Green Supermix solution (Bio-Rad), 1 μL of 5 mM specific primers, and 8 μL of the diluted template. Thermocycling conditions were as follows: 95°C for 3 min followed by 40 cycles of 95°C for 30 s, 60°C for 30 s, and 72°C for 20 s, with a melting curve detected at 95°C for 1 min, 55°C for 1 min, and detection of the denature time from 55°C to 95°C. Each sample was quantified in triplicate and normalized using Ubiquitin10 as the internal control. Gene-specific primer pairs were designed and are available upon request.

### Chromatin Immunoprecipitation (ChIP) assay

ChIP assay was performed as described (Gendrel et al., 2005). Chromatin extracts were prepared from light treated 4-day old seedlings treated with formaldehyde. The chromatin was sheared to an average length of 500 bp by sonication and immunoprecipitated with anti-acetylated histone H3K9K14 (catalog no. 06-599; Millipore). The DNA cross-linked to immunoprecipitated proteins was analyzed by real-time PCR. Relative enrichments of various regions of COP1 over Col-0 were calculated after normalization to Ubiquitin10. Each of the immunoprecipitations was tested and analyzed in triplicates, and each sample was quantified at least three times.

### Protein Extraction and Gel-Blot Analysis

Total protein extraction was conducted as described (Hsieh et al., 2000). Four-day old seedlings of wild type Col-0, all mutant, and overexpression lines were continuously grown under different light treatments and duration. After the treatment, the seedlings were harvested and frozen immediately for protein extraction. One hundred micrograms of total protein were separated on a 12% SDS-polyacrylamide gel, blotted on immuno-blot PVDF (Bio-Rad), and detected using anti-HY5, anti-COP1, and anti-RPN6 specific antibodies.

### Protein-Protein Interaction

For the bifluorescence complementation (BiFC) assay, HDA15 and COP1 coding sequences were cloned into the pCR8/GW/TOPO vector and used for recombination into its corresponding destination vectors, pEG201-N-YFP and pEG202-C-YFP, using LR mix. Kanamycin was used for bacterial selection. Purified plasmids were then analyzed for DNA sequencing for confirmation and further used for protoplast PEG transfection and imaging.

Pull-down assay was carried out as described (Yang et al., 2008) with some modifications. For GST pull-down, GST and GST-HDA15 recombinant proteins were incubated in 30 mL of GST resin with binding buffer (50 mM Tris-Cl, pH 7.5, 100 mM NaCl, 0.25% Triton X-100, and 35 mM β-mercaptoethanol) for two h at 4°C. The binding reaction was washed three times with the binding buffer, and then the COP1-His recombinant protein was added and incubated for an additional two h at 4°C. For His pull-down, His and COP1-His recombinant proteins were incubated in 30 mL of His resin in a phosphate buffer (10 mM Na2HPO4, 10 mM NaH2PO4, 500 mM NaCl, and 10 mM imidazole) for two h at 4°C, the binding reaction was washed three times with the phosphate buffer, and then GST-HDA15 recombinant protein was added and incubated for an additional two h at 4°C. After extensive washing (for at least eight times), the “pulled-down” proteins were eluted by boiling, separated by 10% SDS-PAGE, and detected by western blotting using anti-His, anti-GST, or anti-COP1 specific antibody.

### Bioinformatics analysis

The protein sequences of HY5 and PIF3 were retrieved from the NCBI's protein sequence repository database (www.ncbi.nlm.nih.gov/). Putative protein acetylation sites were predicted by Bayesian based algorithm for protein lysine acetylation (Li et al., 2009).

### Statistical analysis

Statistical analyses were conducted using SigmaStat 11 software. One-way ANOVA using the Holm-Sidak method was performed for multiple pairwise comparisons. Similarly, Dunnette’s method was conducted for multiple comparisons using WT as control. However, Kruskal-Wallis One-way ANOVA on Ranks was used to analyze data that failed to pass the Normality test. Significance testing was set at an α of 0.05 or 0.01. Graphs were presented in mean values with error bars set from its standard deviation.

## RESULTS

### Class II HDA mutant lines display light hyposensitivity

T-DNA SALK lines of Class II histone deacetylases were screened, analyzed, and grown under various light conditions where long hypocotyl phenotypes were initially observed. To determine the maximum phenotypic expression of the mutant lines, plants were exposed to varying light fluence rates, light intensities, and durations. Seeds of knockout lines were grown under white light and dark treatment for seven consecutive days. Class II mutant lines were likewise grown under continuous far-red, red, and blue light at high and low light intensities for five consecutive days. Hypocotyl lengths were measured on the 3rd, 5th, and 7th days (Figure 1).

**Fig. 1.**
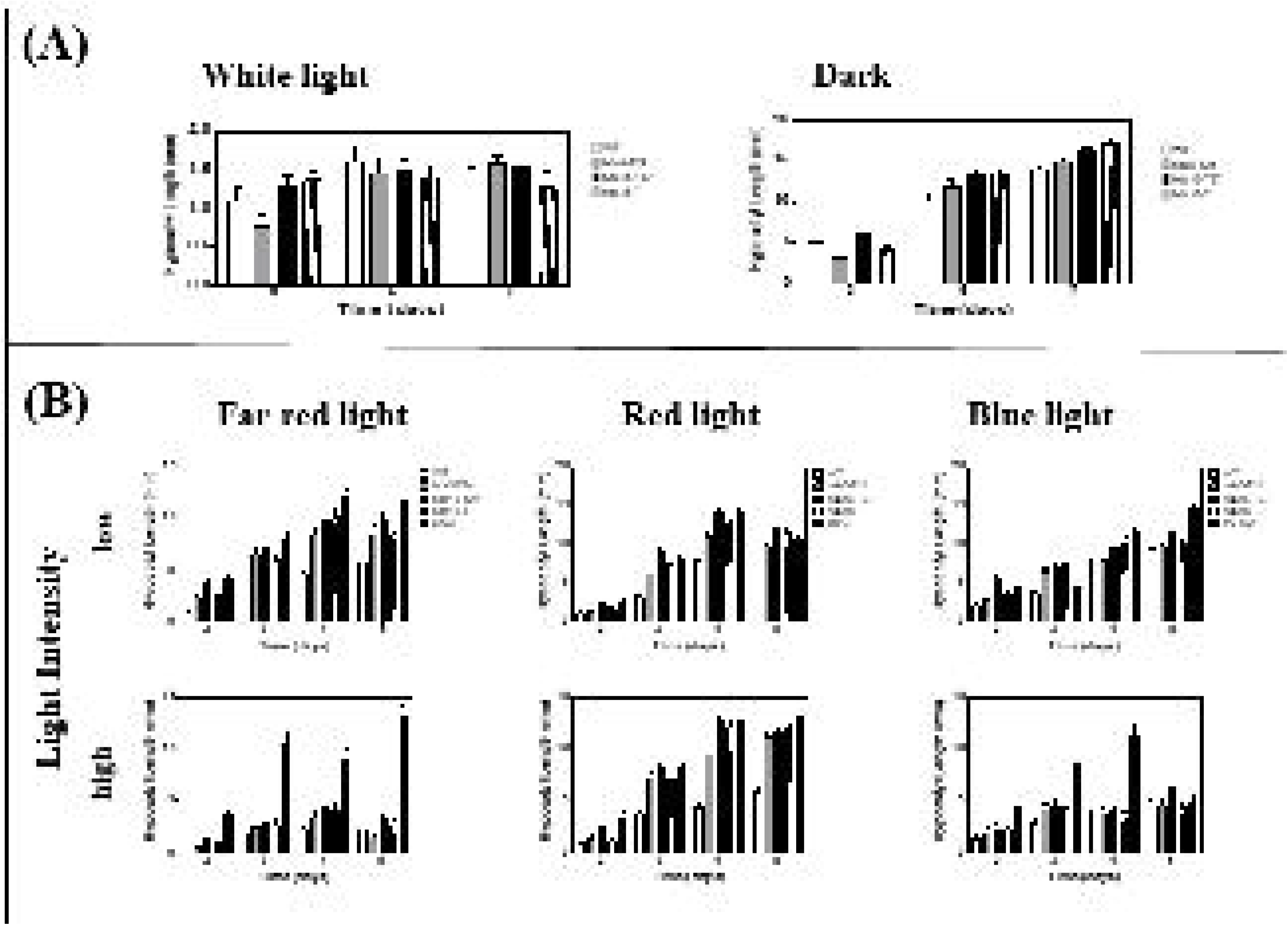
Comparison of light response of *A. thaliana* bearing class II HDA and phytochrome receptor mutations. (A) Photographs and hypocotyl lengths of WT, *hda5*, *hda15,* and *hda18* mutants under to the dark and white light for seven consecutive days. (B) Hypocotyl lengths of the Class II HDA mutants concerning the phytochrome receptor mutants under far-red, red and blue light treatments at different light intensities for seven days. Light intensity treatments were designated as low (L, upper panel) or high (H, lower panel): far-red, H at 18.065 and L at 2.77 μmol m^−2^ s^−1^; red light, H at 22.938 and L at 1.77 μmol m^−2^ s^−1^; and blue light, H at 11.095 and L at 3.84 μ mol m^−2^ s^−1^. Data presented are mean values with error bars set from its standard deviation. Asterisks indicate significant difference, where single asterisk at 95% and double asterisk at 99% level of confidence.

Results indicate that loss-of-function lines of HDA5, HDA15, and HDA18 were generally normal under white light treatment. However, in the absence of light, significantly longer hypocotyls starting on the 3^rd^ day for *hda15-t27* and 4^th^ day for *hda5-t24* and *hda18-t* were observed. The hyper-skotomorphogenic phenotype exhibited by Class II HDA mutants after four days of dark treatment was noticeably pronounced as those displayed by the *phyA* and *phyB* mutants. This behavior suggests that HDA15 and HDA18 may act downstream of the phytochrome signaling pathway.

Furthermore, light hyposensitive phenotypes of T-DNA knockout mutants were significantly prominent at low light intensities after 3 to 4 days of continuous light treatment. Under far-red, red, and blue light treatments, four days of constant light treatment not only induced long hypocotyls but also yielded bigger, more developed open cotyledons with longer petioles indicating that Class II histone deacetylases might act as positive photomorphogenic regulators. Hypocotyl length of mutant lines eventually leveled off with the wildtype on the fifth day of light treatment, indicating that these histone deacetylases could be active at the early stages of growth and development.

All in all, among all the Class II HDAs tested, *hda15-t27* mutants had the most prominent hyposensitive phenotype in both light and dark conditions. Thus, this study focused on elucidating the function of HDA15 in photomorphogenesis.

### HDA15 targets the deacetylation of COP1 promoter

Like the long-hypocotyl phenotype displayed by the *hy5* mutant, we hypothesize that HDA15 acts as a positive regulator of photomorphogenesis. To determine if HDA15 is indeed involved in photomorphogenesis, expression levels of downstream positive regulators of photomorphogenesis were assessed *via* RT-PCR and qPCR. As presented in Figure 2, among the key positive regulators, which are known to be targeted by acetylation upon light exposure, include HY5, LHB1B1, LHCA1, RBCS-1a, and 1b, CAB1, and CAB2 (Guo et al., 2008; Charron et al., 2009). However, most of these genes, including HY5, were significantly reduced in all the light treatments, which explain the long-hypocotyl phenotype suggesting that a negative regulator catapulted the downregulation of these genes, possibly targeted by HDA15.

**Fig. 2.**
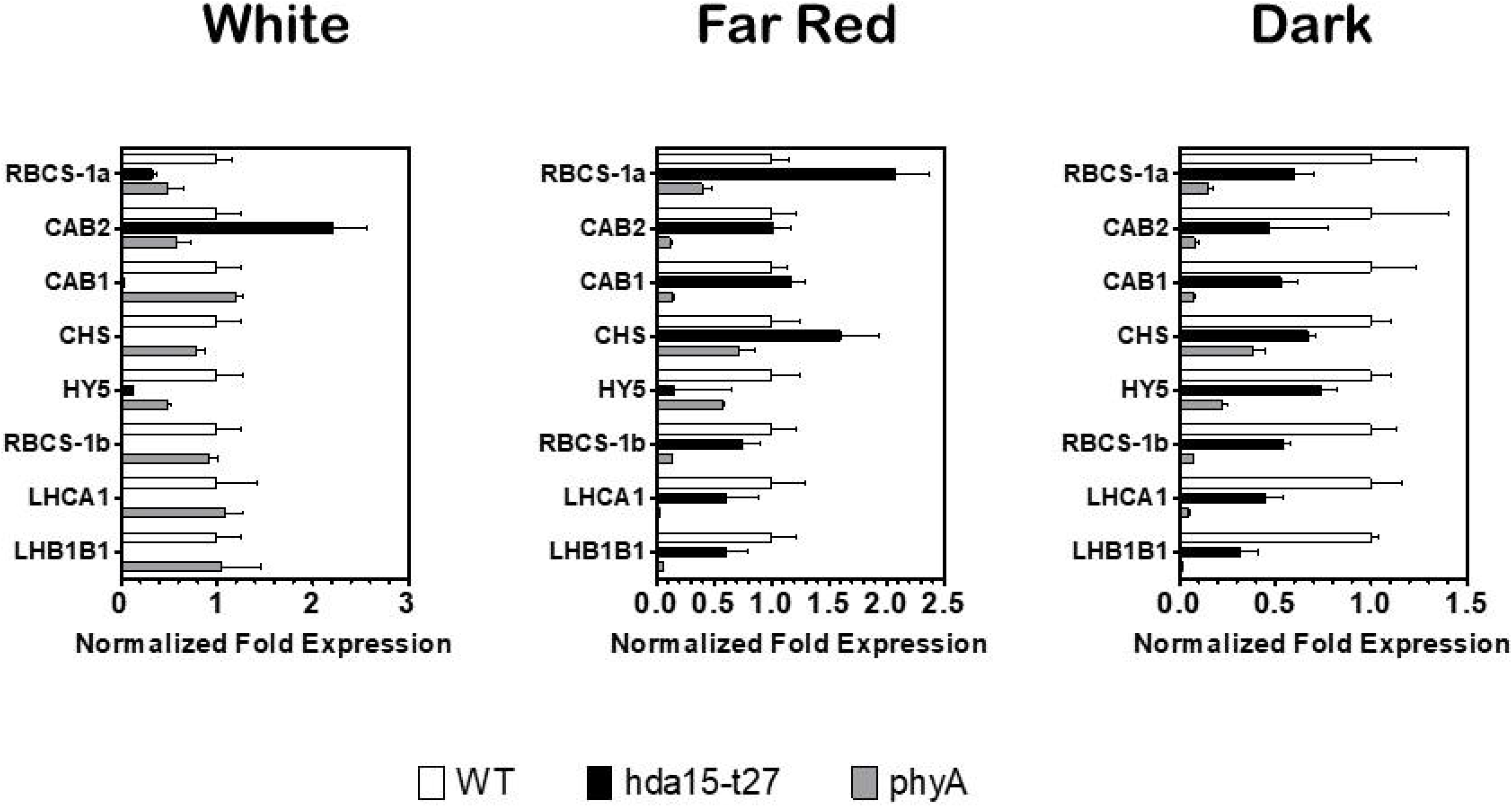
Down-regulation of positive regulators to photomorphogenesis in AtHDA15 knockout lines. mRNA expression levels of positive regulators of photomorphogenesis were assessed in *A. thaliana* exposed to (A) white light, (B) far-red light, and (C) the dark. Expression levels reflected were normalized to ubiquitin.

To further assess the regulatory role of HDA15, mRNA expression profiles of key regulators of photomorphogenesis in *hda15* mutant and overexpression lines were conducted (Figure 3). Compared to white light and far-red light treated plants, dark-treatment has eminently amplified HY5 transcripts levels up to 7-folds in HDA15 overexpression lines. Concomitantly, mRNA levels of COP1 were significantly elevated 3-folds in *hda15* mutants, while its overexpression leads to COP1's extreme downregulation compared to wild type. ChIP assay using anti-H3K9K14ac further reveals that the promoter and start sites of COP1 were highly acetylated in the absence of HDA15 in dark-treated plants. The comparative acetylation profiles suggest that HDA15 targets the deacetylation of COP1 in the dark.

**Fig. 3.**
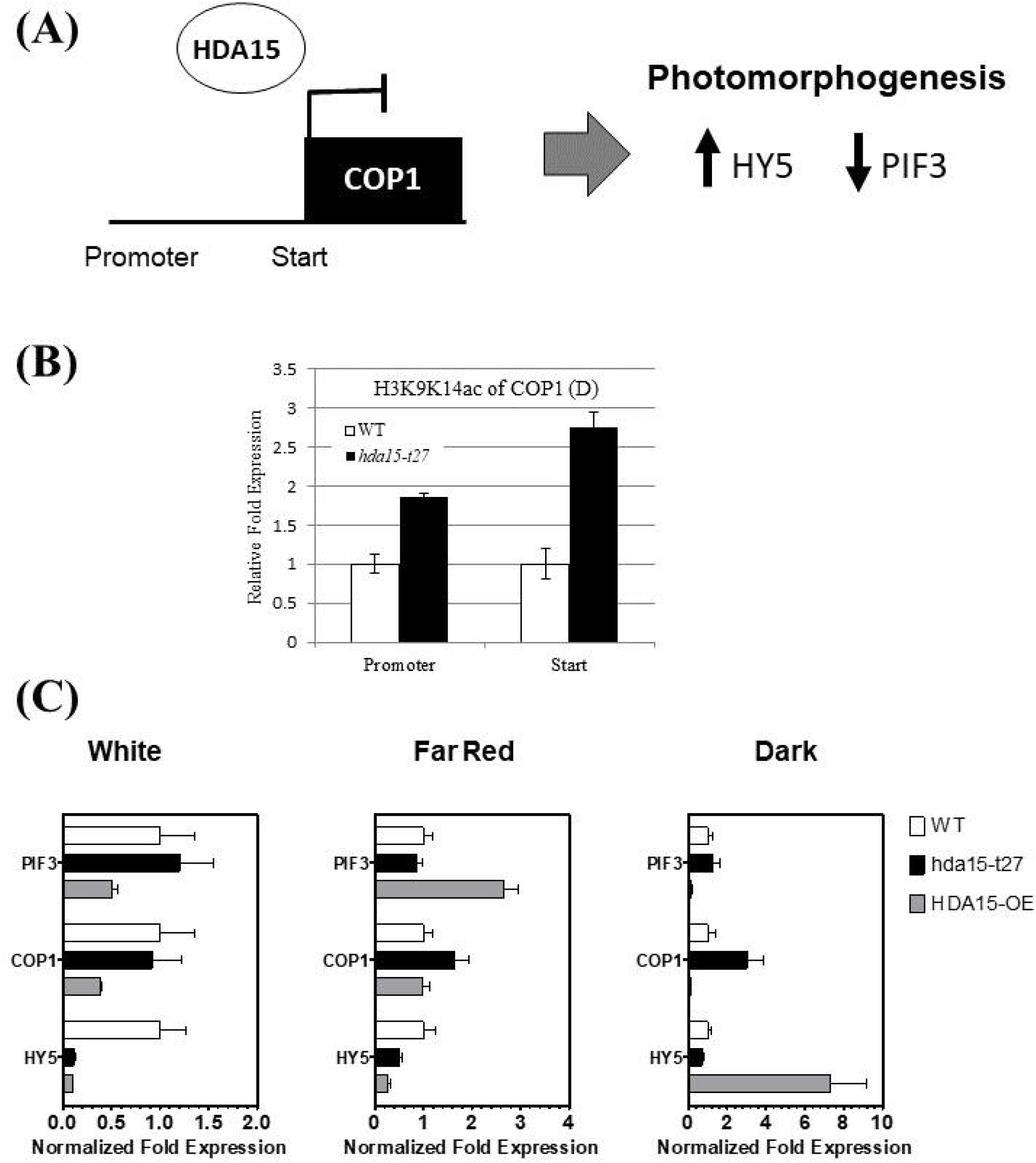
mRNA expression profiles of key regulators of photomorphogenesis in HDA15 mutant and overexpression lines. (A) Schematic diagram of repressive action of HDA15 on COP1 locus. (B) ChIP assay using anti-H3K9K14ac further reveals that the promoter & start sites of COP1 were highly acetylated in the absence of HDA15 in dark-treated plants. (C) Compared to white light and far red light treated plants, dark-treatment has eminently amplified HY5 transcripts in HDA15 overexpression lines. Concomitantly, mRNA levels of COP1 were significantly elevated in *hda15* mutants while its overexpression lead to COP1’s extreme downregulation

### Light signals HDA15 nuclear localization while dark treatment induces cytoplasmic translocation

Hypocotyls of 4-day old HDA15-GFP transformants were used to determine its localization under varying light conditions (Figure 4). Under white light conditions, HDA15 remained to be prominently nuclear. When grown under far-red light, HDA15-GFP concentrated in the nucleus, which is similar to blue and red light treated lines. Consequently, seedlings grown in the dark exhibited strong signals at the cytoplasmic area, although small quantities were still detected at the nucleus. Similar results were observed when HDA15-GFP were transiently expressed in protoplasts under various light treatments.

**Fig. 4.**
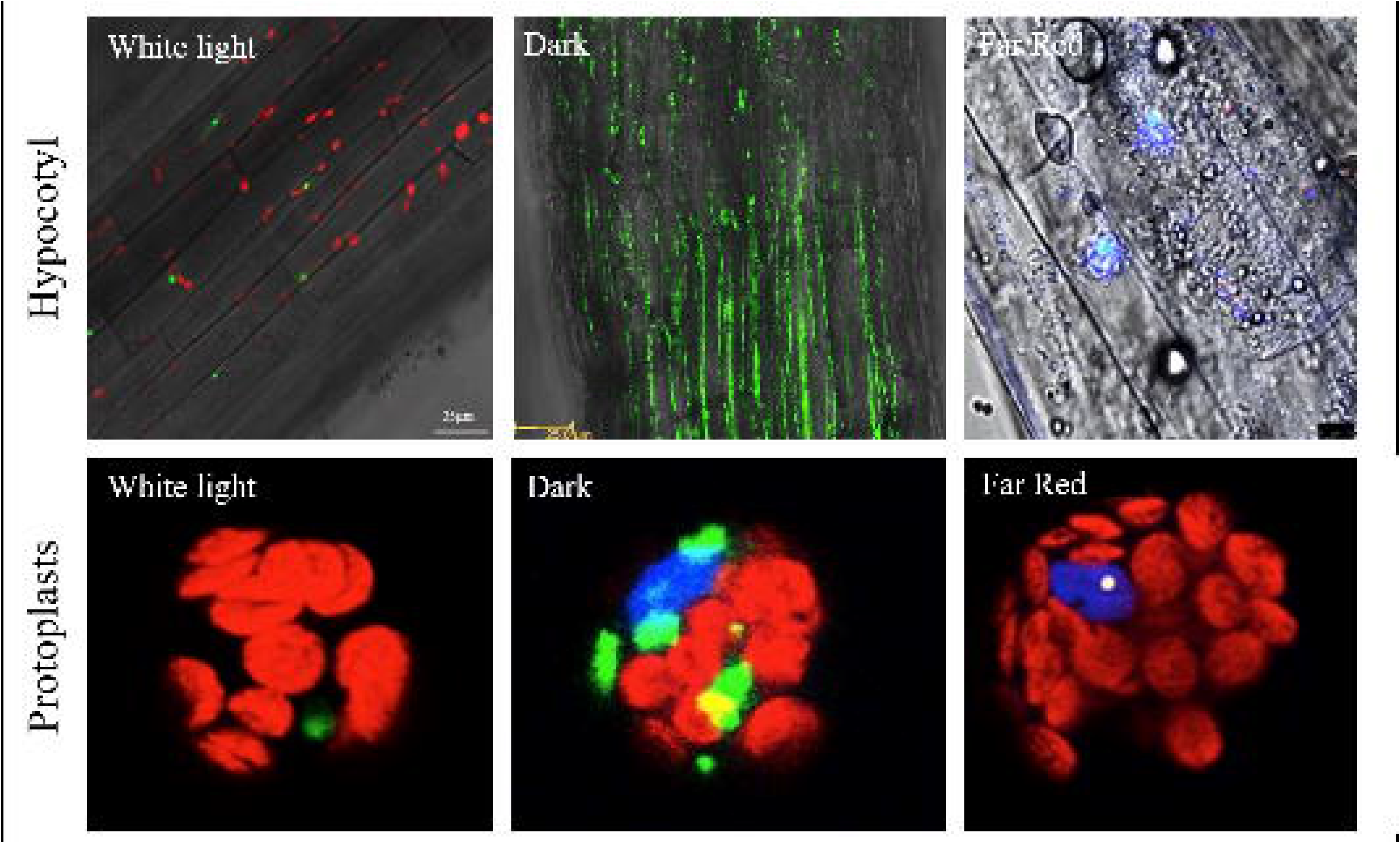
Subcellular localization of AtHDA15 under different light treatments. HDA15-GFP/YFP was localized in the nuclei of hypocotyls and protoplasts of transgenic lines grown under white light and different light treatments. Note: In dark treated plants, HDA15 has a pan-cytoplasmic distribution. Hoechst was used as nuclear marker

As mammalian Class II HDAs undergo nucleocytoplasmic shuttling, plant Class II HDAs HDA15, in particular, likewise exhibit similar functional regulatory mechanisms. This was further illustrated by Alinsug et al. (2012) where HDA15 depends on its own NLS and NES signals for nucleocytoplasmic shuttling. As revealed in their study, HDA15-GFP transfected protoplasts exhibited nuclear localization after 18 hours of white light incubation. Further dark treatment for 3 hours elicited partial cytoplasmic translocation of HDA15-GFP. After one hour, re-exposure of the protoplasts to white light led to its complete nuclear import (Figure 5). The observed response demonstrates that light indeed drives the nucleocytoplasmic shuttling of HDA15.

**Fig. 5.**
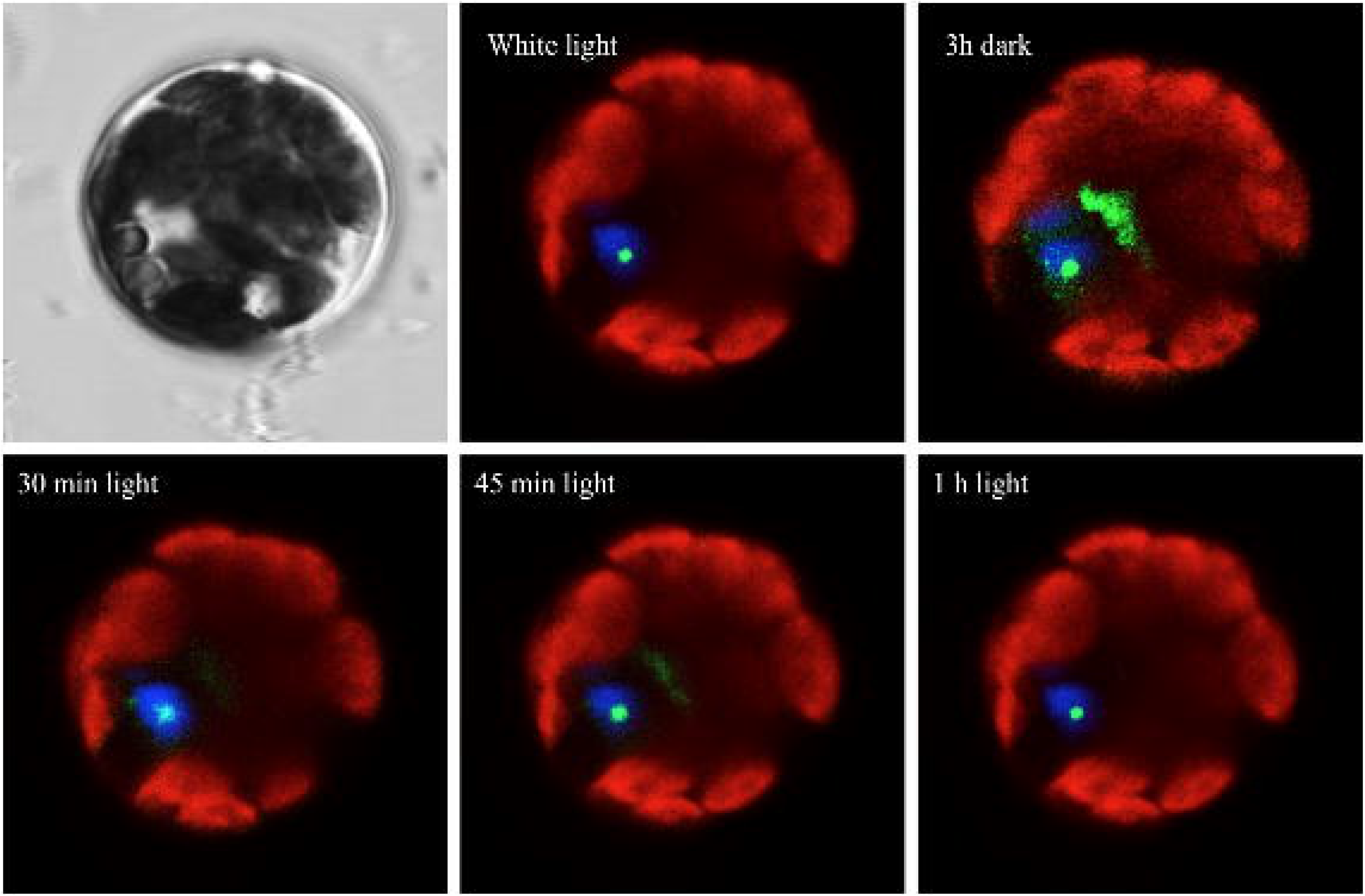
Time-course monitoring of nucleocytoplasmic shuttling and pan-cytoplasmic scattering of HDA15. Transfected protoplasts were incubated under white light overnight then covered with foil for 3 hours and re-exposed to white light. HDA15-GFP transfected protoplasts exhibited nuclear localization after 18h of white light incubation. Further dark treatment for 3h elicited partial cytoplasmic translocation of HDA15-GFP. Re-exposition to white light after 1h lead to its complete nuclear import. VirD2NLS was co-transfected as nuclear marker (blue).

### COP1 triggers the nuclear localization of HDA15

Scanning through the amino acid sequence of HDA15, it contains a type par4 nuclear localization signal (NLS) near the amino end, two overlapping bipartite NLS near the carboxyl end, and a nuclear export signal (NES) near the C terminal half. Our previous study has indicated that these signals navigate the subcellular compartmentalization of HDA15, and that light, in general, drives the nucleocytoplasmic shuttling of HDA15 (Alinsug et al., 2012). Albeit the exact mechanisms controlling its NLS and NES signals upon light exposure remains obscure. Thus, it is speculated that the only way for HDA15 to detect light is through the photoreceptors; their inactivation upon the termination of light may have initiated HDA15’s nuclear export.

To test this hypothesis, HDA15-GFP/YFP was transiently expressed in protoplasts of mutant photoreceptors *phyA, phyB*, and *cry1xcry2*. As shown in Figure 6, all these mutant photoreceptor protoplasts displayed nuclear localization of HDA15 suggesting that the nuclear import of HDA15 is not dependent on light quality nor controlled singly by the phytochrome/cryptochrome signaling cascades, but implicates the involvement of all photoreceptors. The downstream target of all four photoreceptors, COP1, appears to have a significant impact on HDA15's nuclear localization, where *cop1-4* mutant protoplasts exhibited speckled distribution of HDA15 in the cytoplasm immediately near the vicinity of the nucleus. Overall, our results suggested that the activation of photoreceptors did not trigger HDA15 nuclear localization. Instead, the shuttling observed was likely attributed to the presence of COP1.

**Fig. 6.**
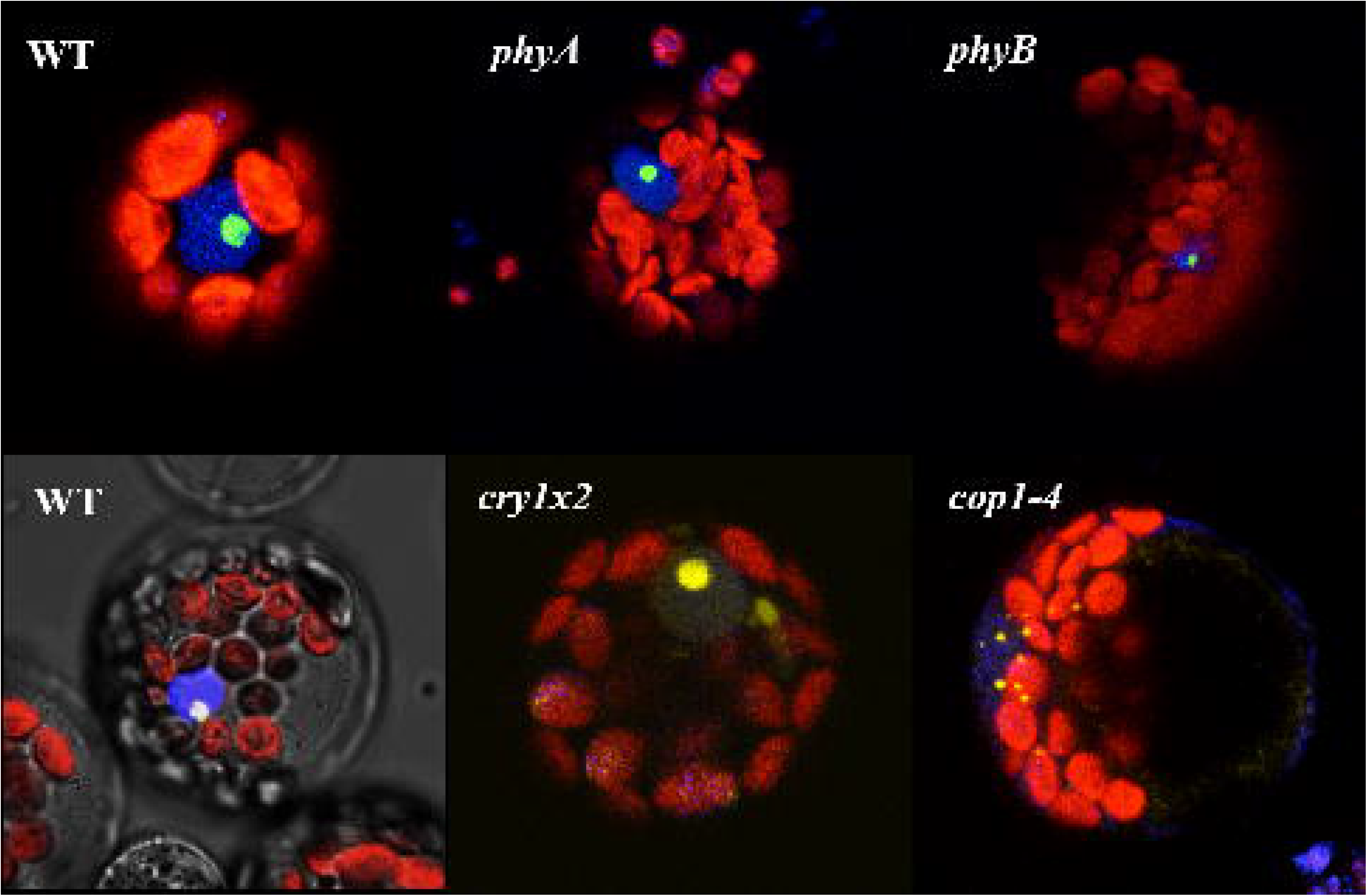
COP1 activates nuclear localization of HDA15. HDA15 localization in protoplasts of mutant photoreceptors were all nuclear. However, when transiently expressed in *cop1-4* mutant protoplasts, HDA15 exhibited cytoplasmic localization indicating that COP1 ultimately influences nuclear localization of HDA15. VirD2-NLS was co-transfected as nuclear marker (blue)

### HDA15 directly interacts with COP1 inside the nucleus

The human Class II HDAC6 has been well studied to catalyze non-histone proteins such as α-tubulin, cortactin, and HSP90, as well as, bind to ubiquitinated proteins inhibiting their proteosomal degradation via its zinc finger ubiquitin-binding protein (ZnF-UBP) domain (Hook et al., 2002; Kovacs et al., 2005; Luxton and Gundersen, 2007; Valenzuela-Fernandez et al., 2008). Although the corresponding plant ortholog of HDAC6 is unclear, HDA15 contains a RanBP zinc finger analogous to ZnF-UBP of HDAC6. To determine if HDA15 can associate with non-histone proteins such as COP1, pull-down and BiFC assays were conducted (Figure 7). Direct binding between HDA15 and COP1 was illustrated via pull-down assay using anti-His, anti-GST, and anti-COP1 antibodies where HDA15 and COP1 were fused with GST- and His-tags, respectively. This interaction was further confirmed *in vivo* using BiFC demonstrating their direct binding inside the nucleus both under white light and dark conditions. Although it has been consistently demonstrated that HDA15 is nuclear under all the light treatments, the abrogation of COP1 in *cop1-4* mutant protoplasts renders it cytoplasmically speckled.

**Fig. 7.**
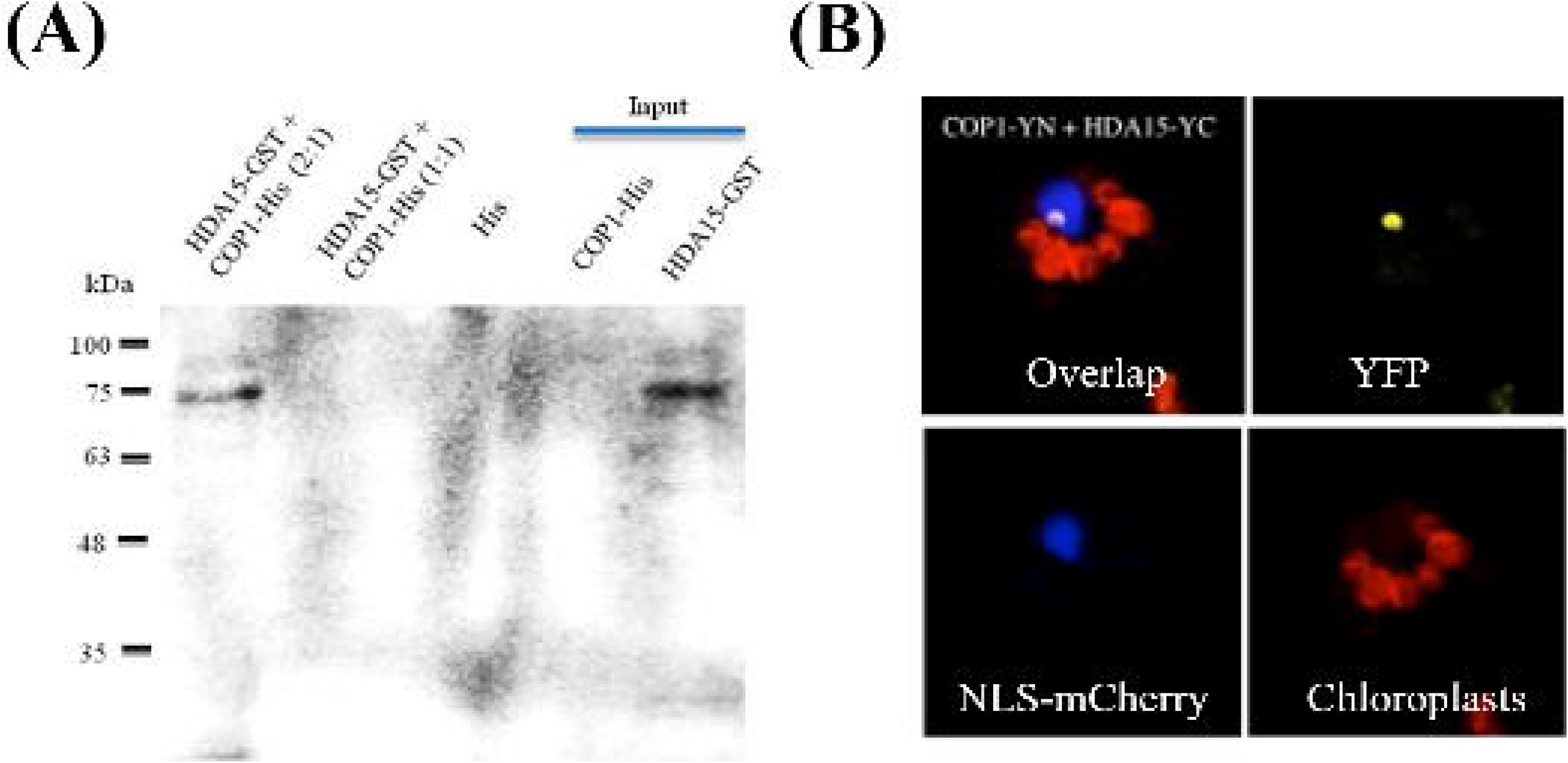
AtHDA15 directly associates with COP1. **(A)** *In vitro* and **(B)** i*n vivo* interaction between COP1 and AtHDA15 indicates their direct binding inside the nucleus. AtHDA15-COP1 interaction was further confirmed via pull down assay using anti-His antibody where HDA15 was fused with GST and COP1 with His tags. Interaction of COP1 and AtHDA15 in isolated protoplasts was visualized 18–24 hr after transfection using bifluorescence complementation assay under laser confocal microscopy. AtHDA15 and COP1 genes were recombined with N-YFP and C-YFP. VirD2-NLS was used as nuclear marker

### HDA15 positively regulates photomorphogenesis by repressing COP1

It has long been hypothesized that COP1 may recruit deacetylases or chromatin remodeling factors that can repress transcription from target promoters (Holm and Deng, 1999). If HDA15 functions as a co-repressor of COP1, then the mutant lines of HDA15 should exhibit *cop1*-like phenotypes. On the contrary, *hda15-t27* lines manifested light hyposensitivity similar to *hy5* mutants with down-regulated levels of HY5 and other positive regulators of photomorphogenesis. Thus, HDA15 may indirectly regulate HY5 by repressing COP1, making it a positive regulator of photomorphogenesis.

To establish the genetic relationship between HDA15 and COP1, we generated the double mutant *cop1-4xhda15-t27*. As shown in Figure 8, the cross between *hda15-t27* and *cop1-4* mutants exhibited short hypocotyl phenotypes under white light and dark treatments and dwarfed phenotypes in 3-week old plants all reminiscent of *cop1-4* mutants. These responses suggest that COP1 is epistatic to HDA15. Therefore, HDA15 targets the repression of COP1 via direct binding, thereby attenuating its capacity to ubiquitinate HY5 and its targets.

**Fig. 8.**
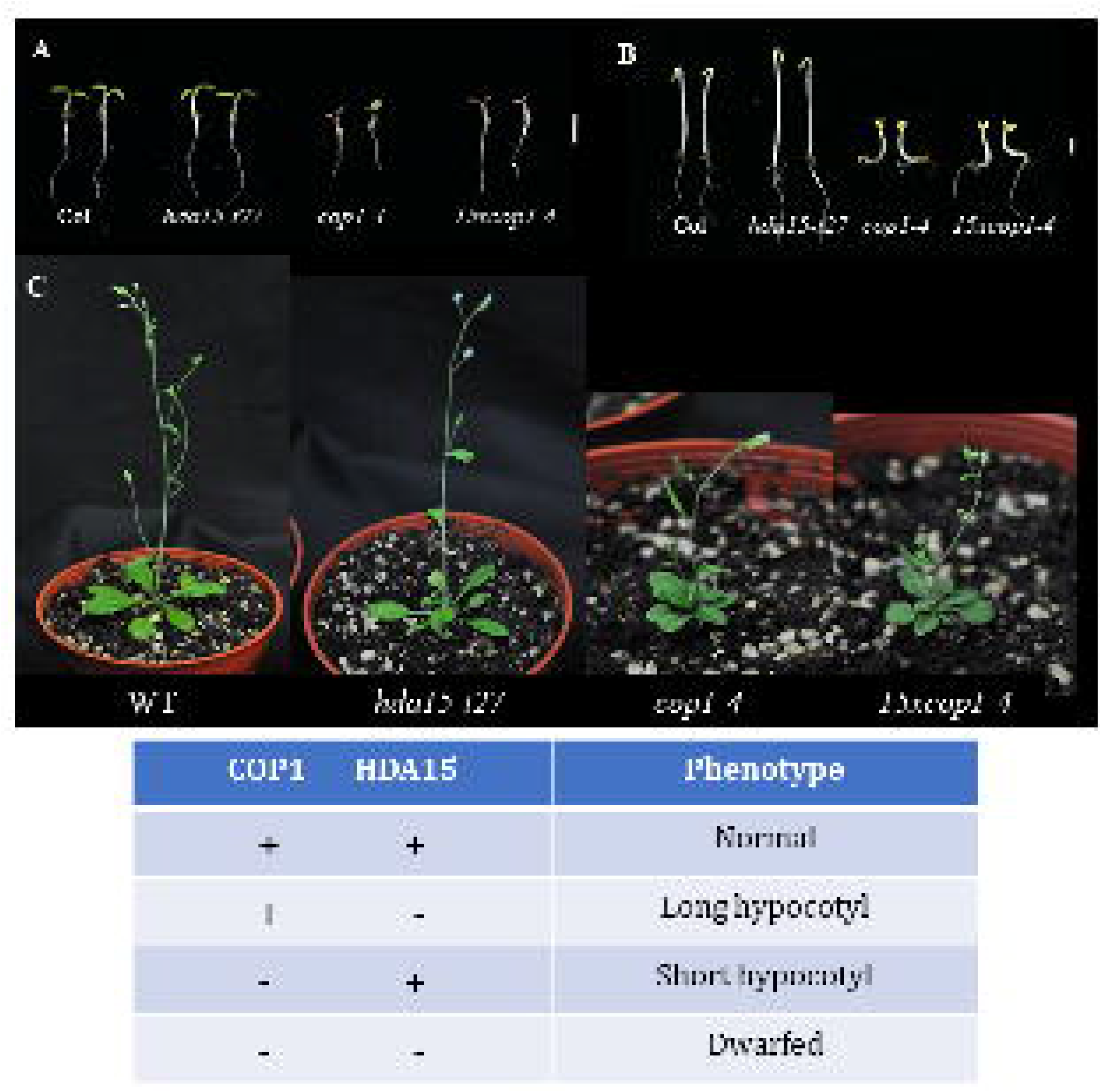
Double mutant *hda15-t27xcop1-4* phenotype. The cross between *hda15-t27* and *cop1-4* mutants exhibited short hypocotyl phenotypes under white light (A) and dark (B) treatment for four days and dwarfed phenotypes in 3-week old plants (C) which are reminiscent of *cop1* mutants. This indicates that COP1 is epistatic to HDA15

### Ubiquitinated sites of HY5 and COP1 are less stable than HDA15

Considering that COP1 is an E3 ligase, the binding of HDA15 with COP1 raises pertinent questions as to whether COP1 ubiquitinates HDA15 or HDA15 represses COP1. Aside from the double mutant cross between *had15* and *cop1* resulting to *cop1*-like phenotype, bioinformatics analysis using three algorithms, namely: Ub-Pred, BDM-PUB, and UbiSite, were used to predict ubiquitination sites comparing HY5, COP1, and HDA15. Based on the algorithms' consensus, as presented in Table 1, HY5 was found to contain five residues that are actively targeted for ubiquitination, rendering it the most unstable among the three, with instability index of 68.21. COP1 follows HY5 with an instability index of 47.13, with four consensus residues prone to ubiquitination. On the other hand, it appears that HDA15 is the most stable among the three proteins with an instability index of 37.11, noting that it only has two residues as prospective sites for ubiquitination. This bioinformatics analysis hence reaffirms our prior notion that HY5 is targeted for ubiquitination by COP1 in the dark during skotomorphogenesis. With HDA15 being more stable than COP1, the direct interaction between HDA15 and COP1 leads us to conclude that HDA15 represses COP1.

**Table I.** Predicted ubiquitinated residues in AtHDA15, COP1 and HY5 based on three algorithms.

| Gene | ID | Length<br>(aa.) | Ub-Pred* | BDM-PUB** | UbiSite*** | Consensus | Instability<br>Index |
| --- | --- | --- | --- | --- | --- | --- | --- |
| AtHDA15 | Q8GXJ1 | 552 | 14 (0.75); 129 (0.65); 139 (0.80); 167 (0.69) | 14 (2.34); 46 (2.87); 113 (0.31); 129 (0.76); 139 (0.59); 265 (0.90); 305 (1.25); 306 (1.96); 375 (1.26); 435 (2.14); 458 (1.49); 503 (2.08); 516 (0.48); 517 (0.54); 521 (0.88); 523 (1.02); 552 (4.49) | 46 (0.55); 129 (0.38); 139 (0.45); 167 (0.62); 205 (0.40); 265 (0.58); 305 (0.67); 306 (0.53); 328 (0.46); 375 (0.31); 458 (0.35); 503 (0.33); 532 (0.25); 536 (0.30); 537 (0.35) | 129; 139 | 37.11<br>(stable) |
| COP1 | P43254 | 675 | 14 (0.80); 193 (0.66); 273 (0.84); 274 (0.96); 312 (0.74); 328 (0.76); 433 (0.66) | 14 (1.51); 106 (2.66); 109 (0.96); 110 (1.17); 118 (1.86); 139 (1.35); 152 (1.87); 154 (0.70); 176 (0.56); 178 (1.40); 216 (1.21); 263 (2.02); 273 (1.89); 274 (2.79); 295 (0.38); 296 (0.92); 323 (0.91); 328 (1.76); 396 (1.90); 512 (1.04); 549 (1.10); 550 (0.73); 556 (1.65); 653 (0.73); 669 (1.04) | 14 (0.47); 106 (0.47); 109 (0.34); 110 (0.28); 118 (0.62); 152 (0.43); 154 (0.40); 176 (0.39); 178 (0.33); 193 (0.23); 216 (0.44); 234 (0.29); 258 (0.56); 259 (0.47); 263 (0.23); 273 (0.50); 274 (0.51); 312 (0.26); 323 (0.36); 328 (0.28); 396 (0.20); 422 (0.21); 464 (0.48); 512 (0.30); 549 (0.23); 550 (0.45); 556 (0.27); 578 (0.50); 593 (0.56); 619 (0.28); 669 (0.35) | 14; 273; 274;<br>328 | 47.13<br>(unstable) |
| HY5 | O24646 | 168 | 31 (0.95); 53 (0.83); 79 (0.68); 110 (0.70); 121 (0.80); 126 (0.87) | 31 (0.86); 53 (1.95); 79 (1.32); 87 (3.02); 90 (2.34); 93 (1.64); 109 (1.65); 110 (1.05); 126 (0.87); 155 (1.58) | 31 (0.38); 53 (0.43); 79 (0.24); 90 (0.38); 109 (0.33); 110 (0.38); 121 (0.42); 126 (0.46); 149 (0.28); 155 (0.38) | 31; 53; 79; 110;<br>126 | 68.21<br>(unstable) |

\* green (low confidence): score range ( $0.62 \leq s \leq 0.69$ ), sensitivity (0.464), specificity (0.903); blue (medium confidence): score range ( $0.69 \leq s \leq 0.84$ ), sensitivity (0.346), specificity (0.950); red (high confidence): score range ( $0.84 \leq s \leq 1.00$ ), sensitivity (0.197), specificity (0.989)
\*\* Threshold= 0.3
\*\*\* Low threshold; green (low confidence level): score range ( $s \leq 0.50$ ), sensitivity (0.8648), specificity (0.3396); red (high confidence level): score range ( $0.75 \leq s$ ), sensitivity (0.3754), specificity (0.8628)

## DISCUSSION

Histone active marks such as H3K9K27ac signals transcriptional activation whereby light induces the acetylation of positive photomorphogenic regulators functioning as chlorophyll a/b-binding protein, peroxidase, citrate synthase, RBCS small chain 1B, ribosomal protein, PSII-light-harvesting complex, HYH, HY5 and its downstream targets CHS, IAA3, RBCS-1a during photomorphogenesis (Benhamed et al., 2006; Guo et al., 2008; Charron et al., 2009). On the other hand, histone deacetylation stimulates chromatin compaction triggering transcriptional quiescence. Among the RPD3/HDA1-like superfamily of HDAs, Class I histone deacetylases HDA6 and HDA19 display negative regulatory roles in photomorphogenesis. Together with PHYB, Tessadori *et al.* have identified HDA6 as a regulator of light-induced chromatin condensation (Tessadori et al., 2009). Similarly, HDA19 is suspected of repressing many light-regulated genes by deacetylating H3K9K14 at the transcriptional start site of the phyA locus (Benhamed et al., 2006; Jang et al., 2011).

On the other hand, it appears that Class II HDAs play a positive regulatory role in the light signaling network with HDA15 repressing COP1 at its locus and modulating its activities via sub-compartmentalization and direct interaction. As HDA5 and HDA18 also exhibit long-hypocotyl phenotypes at varying light treatments, it remains a conundrum of how these and other Class II HDAs function during photomorphogenesis. These findings fine-tune our prior understanding of the role of histone modifications in light-regulated gene expression. Light not only induces the acetylation of key positive regulators of photomorphogenesis but also renders the simultaneous repression of vital negative regulators such as COP1 via direct interaction. Taken together, our results indicated that histone acetyltransferases and deacetylases act as multi-functional and potent chromatin-modifying enzymes during photomorphogenesis.

While other mutant Class II HDAs also manifest light hyposensitivity, only HDA15 is driven by light to shuttle in and out of the nucleus. The environmental cue or stress that would signal the nuclear import of the other Class II HDAs and their functional characterization remain to be investigated as most of them have been observed to be cytoplasmic (Alinsug et al., 2012; Tran et al., 2012).

Histone deacetylases constitute a family of nuclear enzymes known primarily for their role in transcriptional repression. Class II HDAs, in particular, are known to be versatile regulators catalyzing the deacetylation of both histones and non-histone proteins. The mammalian HDAC6 modulates a wide variety of cellular activities involving complex interactions with α-tubulin, cortactin, HSP90, and other proteins (Matthias et al., 2008; Valenzuela-Fernandez et al., 2008). Similarly, HDAC1 complexes with MDM2, an E3 ubiquitin ligase, deacetylate p53, and promote its ubiquitination (Ito et al., 2002). In the case of AtHDA15, it attenuates COP1 modulating its repressive activities via direct interaction, subcellular compartmentalization, and chromatin compaction at its transcriptional start site. Based on our results, this versatile regulator not only functions as a traditional histone deacetylase silencing key regulators of photomorphogenesis, specifically COP1, but it also directly interacts with COP1 likely attenuating its E3 ligase activities.

The specific molecular mechanisms for the degradation and/or cytoplasmic export of COP1 in the nucleus under light conditions remain obscure. The nuclear interaction of HDA15 with COP1 under light conditions may support the notion that COP1's depletion may be enhanced by the binding of HDA15 to induce auto-ubiquitination. Based on studies conducted by Alinsug *et al*. (2020), overexpression of HDA15 significantly reduced COP1 protein levels and elevated HY5 protein concentrations in various light treatments. However, *hda15* mutant lines did not incur any changes in its COP1 protein levels. Thus, biochemical studies elucidating on the possibility of induced auto-ubiquitination of COP1 via HDA15 binding would be an interesting research path to pursue.

In the presence of light, COP1 is mainly cytoplasmic, although some trace amounts are left inside the nucleus. This purges HDA15 to shuttle into the nucleus binding to COP1 to attenuate its E3 ligase activity against HY5 regulating photomorphogenesis. In the dark, contrarily, COP1 exclusively localizes inside the nucleus and targets the ubiquitination of HY5 to advance skotomorphogenesis. While the majority of HDA15 translocates into the cytoplasm, partial amounts left in the nucleus target the deacetylation of the transcriptional start site of COP1 locus and/or interact with COP1 to modulate its repressive activities. The molecular switches in apportioning the exact amounts of HDA15 to remain active inside the nucleus while others are deactivated inducing their nuclear export remains elusive and requires further investigation.

Presented in Figure 9 is a working model depicting the sub-compartmentalization and modularized activities of COP1 and HDA15 under light and dark conditions. As light signals the nucleocytoplasmic shuttling of HDA15, its nuclear import is primarily dependent on the presence of COP1. Based on our findings, transient expression of HDA15-GFP using *cop1-4* mutant protoplasts exhibits cytoplasmic localization whether it is treated under white light or dark. Since HDA15 have contrasting localization with COP1 in the presence and absence of light, we hypothesize that a concentration gradient on either of the proteins influences the movement of the other in opposing directions, assuming these enzymes are not degraded; or, if a dominant ligand induces the degradation of the other. Thus, further studies are required to unravel the molecular underpinnings of the opposing nucleocytoplasmic translocation of HDA15 & COP1 upon light and dark signaling cues.

**Fig. 9.**
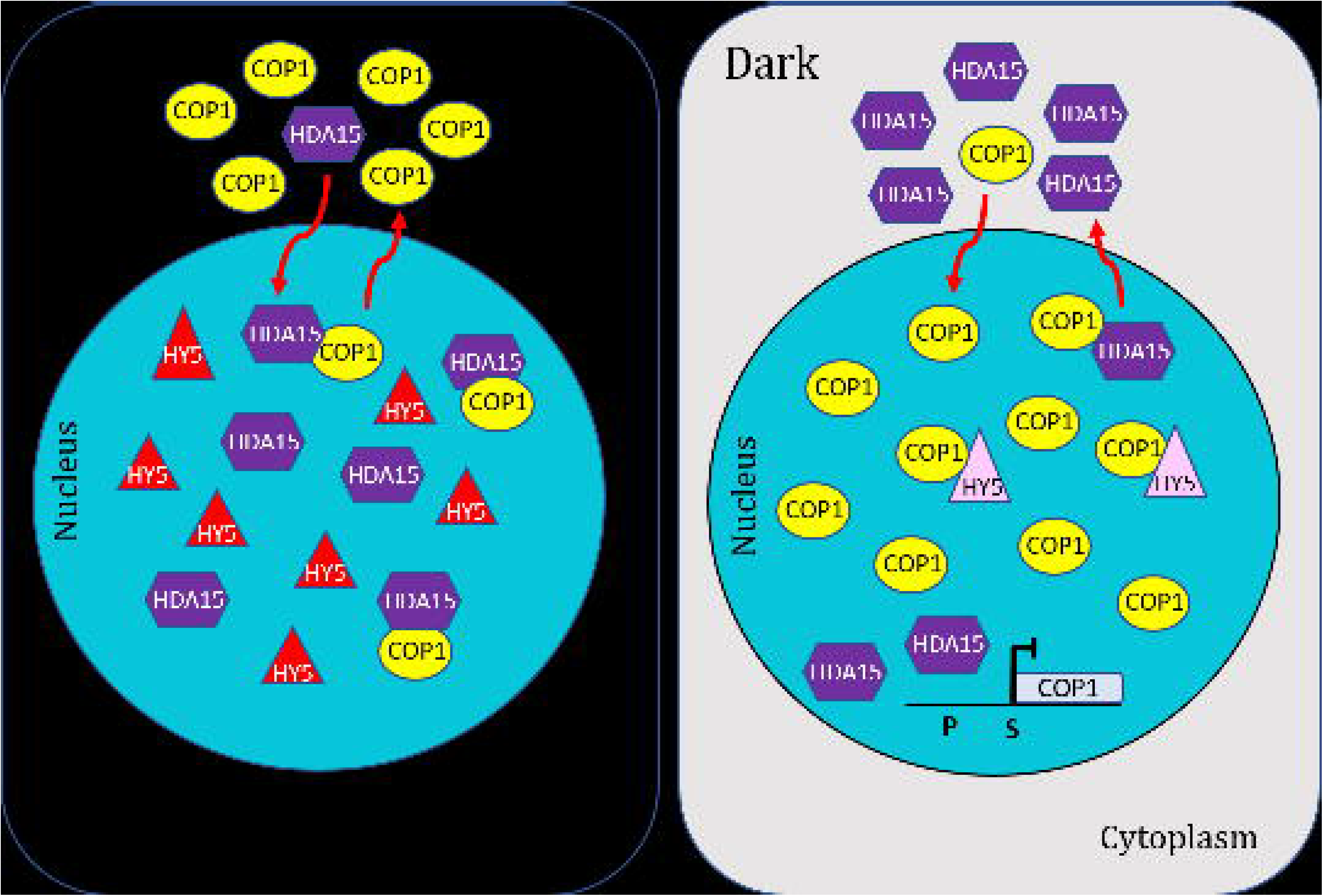
Working hypothesis illustrating modular activities and sub-compartmentalization of COP1 and HDA15 under light and dark conditions. The nucleocytoplasmic movement of HDA15 follows a concentration gradient of COP1 in opposing directions. In the presence of light, COP1 translocates into the cytoplasm purging HDA15 to be nuclear. With some amounts left inside the nucleus, HDA15 binds to COP1 attenuating its E3 ligase activity. In the dark, however, COP1 exclusively localizes inside the nucleus targeting the ubiquitination of HY5 proceeding to skotomorphogenesis. This forces the majority of HDA15 to shuttle into the cytoplasm with some amounts left in the nucleus targeting deacetylation of COP1 start site for transcriptional repression

We also attempted to determine the binding sites of the two repressors by generating different truncated constructs of COP1 and HDA15 containing various combinations of potential protein-binding domains. Surprisingly, only the full-length HDA15 was able to bind the full-length COP1 suggesting strict, proper protein conformation for such interaction.

Previous studies on the conserved domains of RPD3/HDA1-like superfamily show that Zn finger domains are imprinted on several Class II HDAs (Pandey et al., 2002; Alinsug et al., 2009). The monocot ortholog of AtHDA15 in maize, ZmHDA110, has a C3HC4 type RING Zn finger that binds with ubiquitin while the ZnF-UBP of the mammalian Class II HDAC6 binds to ubiquitin specifically at the C termini diglycine motif catalyzed by a deubiquitinase, ataxin-3 (Ouyang et al., 2012). RanBP-ZnF in HDA15 may similarly interact with ubiquitin, such as those found in Npl4 (Alam et al., 2004). Based on sequence analyses on Zn fingers conducted by Wang *et al.*, the molecules have highly conserved cysteine residues and other key motifs in binding ubiquitin (Wang et al., 2003). As illustrated in Supplemental Figure 1, sequence comparison between HDA15's RanBP2-ZnF to the human Np14 (NZF) reveals the presence of the two canonical highly-conserved cysteine residues and a key ubiquitin-binding motif, Thr_95_. Considering the RING-ZnF domain of COP1 catalyzes the transfer of ubiquitin units from E2 transferases to target proteins such as HY5, we speculate that ubiquitin may constitute a recognition site for HDA15 to identify COP1 as an active E3 ligase.

Nevertheless, unlike HDAC6, which binds and transports ubiquitinated proteins to aggresomes, the binding of HDA15 to COP1 undertakes a regulatory role via the attenuation of the latter's E3 ligase activity. Based on bioinformatics analysis comparing the ubiquitination sites of HY5, COP1, and HDA15, it is predicted that HDA15 is more stable than COP1. As expected, HY5 is the least stable making it more prone to ubiquitination. Still, questions remain on the exact nature of binding of HDA15 to COP1 as well as the properties of the molecular switches downstream of the HDA15-COP1 complex.

## Acknowledgments

The authors are grateful to the unwavering support of the various laboratory professors, mentors, colleagues, and staff for the conduct of this research. Special mention goes to Prof. Keqiang Wu and Prof. Hsu-Liang Hsieh of NTU and Prof. Jeong Joo Cheong of SNU for their guidance and access to their facilities. MVA is equally indebted to the scholarship and research fellowship grants funded by the Democratic Pacific Union, National Science Council of Taiwan, National Taiwan University, and Seoul National University.

## Declarations

### Data Availability

The data supporting the findings of this study are available in the paper and from the corresponding author, Malona V. Alinsug, upon request.

### Code Availability

NA

### Authors’ Contribution

All authors contributed to the study conception and design. Material preparation, data collection and analysis were performed by Malona V. Alinsug and Custer C. Deocaris. The first draft of the manuscript was written by Malona V. Alinsug and Custer C. Deocaris commented on previous versions of the manuscript. Both authors read and approved the final manuscript.

### Ethics Approval

NA

### Consent to Participate

NA

### Consent to Publish

All authors read and approved the final manuscript.

---

**Supplemental Fig. 1** Sequence comparison between Zn fingers in human Npl4 (NZF), wild type RanBP2, which has minimal Ubq binding, and RanBP2 in AtHDA15. Conserved residues in NZF were defined by Wang *et al*. (2003), where blue letters indicate semi-conserved residues, green cysteine residues are highly conserved and contact Ubq, and orange residues as key motifs in binding Ubq.

