## Supplementary figures and images for "AtHDA15 attenuates COP1 *via* transcriptional quiescence, direct binding, and sub-compartmentalization during photomorphogenesis"

### Supplemental Figure 1

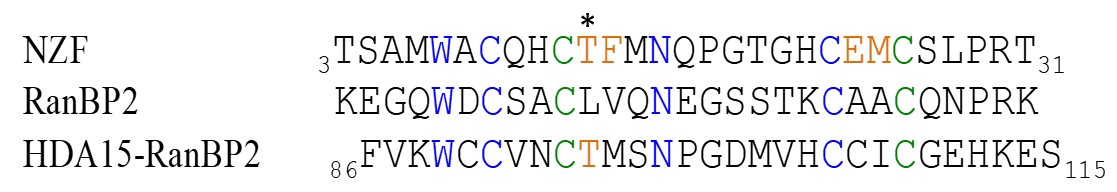
